# Epithelial-stromal cell interactions and ECM mechanics drive the formation of airway-mimetic tubular morphology in lung organoids

**DOI:** 10.1101/2020.12.03.408815

**Authors:** Tankut G. Guney, Alfonso Muinelo Herranz, Sharon Mumby, Iain E. Dunlop, Ian M. Adcock

## Abstract

The complex cellular organisation of the human airway tract where interaction between epithelial and stromal lineages and the extracellular matrix (ECM) make it a difficult organ to study *in vitro*. Current *in vitro* lung models focus on modelling the lung epithelium such as air-liquid interface (ALI) cultures and bronchospheres, do not model the complex morphology and the cell-ECM interaction seen *in vivo*. Models that include stromal populations often separate them via a semipermeable barrier, which precludes the effect of cell-cell interaction or do not include the ECM or the effect of ECM mechanics such as viscoelasticity and stiffness. Here we investigated the effect of stromal cells on basal epithelial cell-derived bronchosphere structure and function through a triple culture of bronchial epithelial, lung fibroblast and airway smooth muscle cells. Epithelial-stromal cross talk enabled formation of epithelial cell-driven branching tubules consisting of luminal epithelial cells surrounded by stromal cells termed bronchotubules. Addition of agarose to the Matrigel scaffold (Agrigel) created a mechanically tunable ECM, where viscoelasticity and stiffness could be altered to enable long term tubule survival. Bronchotubule models enable the investigation of how epithelial-stromal cell and cell-ECM communication drive tissue patterning, repair and development of disease.

**Significance Statement:** Current models of airways diseases such as asthma and COPD do not reflect the physical characteristics of the diseased airway which may impact upon our understanding of disease pathophysiology. We have utilised the physical properties of agarose to modify the 3D stiffness of Matrigel to resemble the human airway. Using a primary airway epithelial cell-derived organoid model we demonstrate that a combined Matrigel/agrigel matrix allows sustained 3D organoid structure and the creation of tubules that can contract in response to a clinically relevant bronchoconstrictor. A complex 3D organoid composed of functioning epithelial cells, smooth muscle cells and fibroblasts may provide opportunities for refined drug discovery programmes.

**Graphical Abstract:** 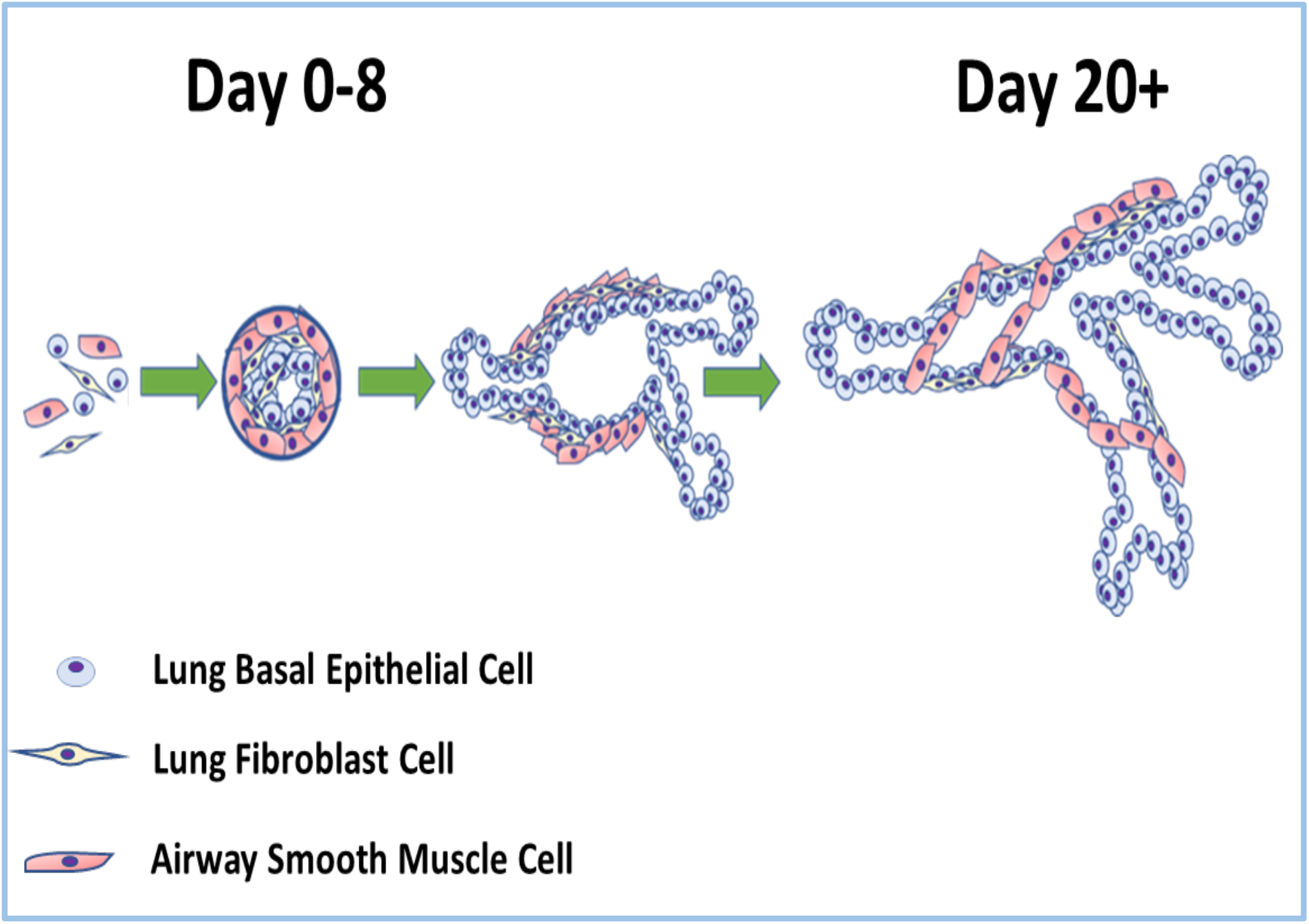

- Mixture of healthy lung basal epithelial cells and healthy lung fibroblast cultured in matrigel result in tubules that fail in 4 days.
- Addition of healthy airway smooth muscle allows for a contractile phenotype.
- Triple culture of cells in a stiffer scaffold agrigel allows maintenance of tubular organoids for a minimum of 20 days.

## Introduction

The airway is a complex organ consisting of highly branched tubes that provide air to the alveoli enabling gas exchange. These tubes contain multiple layers of interacting cells and the development, maintenance and regeneration of the airway is governed by the cell-cell and cell-extracellular matrix (ECM) interactions. Furthermore, the mechanical properties of the ECM in the lower airways are integral to tissue architecture and cellular differentiation [1–3].

The airway is formed from a pseudostratified epithelial layer of mucous and ciliated cells underlined by basal cells attached to a basement membrane ECM [4, 5]. The basement membrane separates the epithelium from the interstitial matrix (lamina propria) that contains immune and fibroblast cells as well as vasculature [4, 5]. Fibroblasts provide structural support by secreting ECM as well as being a main player during cell and ECM repair [6]. Bands of airway smooth muscle form around the lamina propria and control the size of the airway lumen [7]. Perturbation of cell-cell and cell-ECM mechanical dynamics is associated with diseases such as interstitial pulmonary fibrosis (IPF) or chronic obstructive pulmonary disease (COPD) [1–3, 8, 9].

Monolayer cultures of airway epithelium are ineffective in modelling disease reflecting the differences by which primary human cells interact with each other and how they communicate within the airway 3D structure [10, 11]. As a result, more complex models including lung-on-a-chip, have been developed [10, 12, 13], that provide an opportunity to expose cells to environmental stimuli and/or examine the impact of immune cells[12]. However, these 2D models are not representative of the *in vivo* airway environment where neighbouring cells physically interact with each to form tissue within a 3D space [12, 14].

To address this gap 3D organoids have been developed using stem or stem-like cells that self-organise and auto-differentiate in the presence of an ECM scaffold to form the 3D functional multilineage architecture of the originating organ tissue [15–18]. Lung organoids or organospheres form from the aggregation and differentiation of human bronchial or tracheal basal cells that give rise to multiple cell lineages including ciliated and goblet cells [17, 19, 20].

Lung organoids were originally derived from mouse tracheal basal cells, whilst later studies used human bronchial basal epithelial cells to demonstrate that Notch2 and IL-13 inhibition enhanced the proportion of *Foxj1* positive ciliated cells at the expense of mucous producing cells [17].

The use of human pluripotent stem cells (hSPC) has enabled the recapitulation of the tubular architecture of the airways. These lung bronchial organoids (LBO) form in ~170 days and recapitulate the features of the embryonic lung showing alveolar ATI and ATII distal markers distally and the club and mucous cell markers SCGB3A2 and MUC5AC proximally [21]. However, these models do not contain mesenchymal populations such as fibroblasts or airway smooth muscle (ASM) cells that are crucial for the development and maintenance of the airway, for airway contraction and remodelling and are dysregulated in airway diseases such as asthma, COPD and IPF [22–24]. In addition, the effects of the extracellular matrix (ECM) released by mesenchymal cells need to be considered as a key parameter in organoid design, since ECM stiffness profoundly affects cellular behaviour, differentiation and overall organ morphology [25–27].

The critical role of matrix makes organoids an ideal tool to simulate cell-cell interaction *in vitro* as cells can interact freely within an ECM and can simultaneously form contacts with neighbouring cells [28]. In this study, we modelled the effect of normal healthy lung fibroblast and airway smooth muscle (NHLF and NHASM) stromal cells on the development of healthy primary human bronchial epithelial (NHBE) cells. We hypothesised that interactions between NHBE:NHLF:NHASM interaction would change the morphology of epithelial organoid spheroids and aimed to investigate whether co-culture in organoids could mimic a human airway.

## Results

### Branched tubular structures formed by fibroblast (NHLF) and epithelial cell (NHBE) co-cultures

We confirmed previous reports of the formation of spheroidal lumen-containing structures [20, 29](**Figure 1**) and investigated how incorporating fibroblasts into airways epithelial cell culture influences morphology. NHLF and NHBE were first co-cultured. NHLF were seeded as a monolayer and overlayed with 25% Matrigel for 24hr after which NHBE in a 5% Matrigel layer was added. At low NHLF levels (<150,000 cell/ml), NHBE aggregated into spheroidal structures after 0-3 days of culture being similar to those observed in NHBE-only controls as described previously **(Figure 1)**. However, at higher NHLF numbers (>150,000 cell/ml), NHBEs formed rod-like structures that connected distinct epithelial spheres at around 3 days **(Figure 2)**.

**Figure 1.**
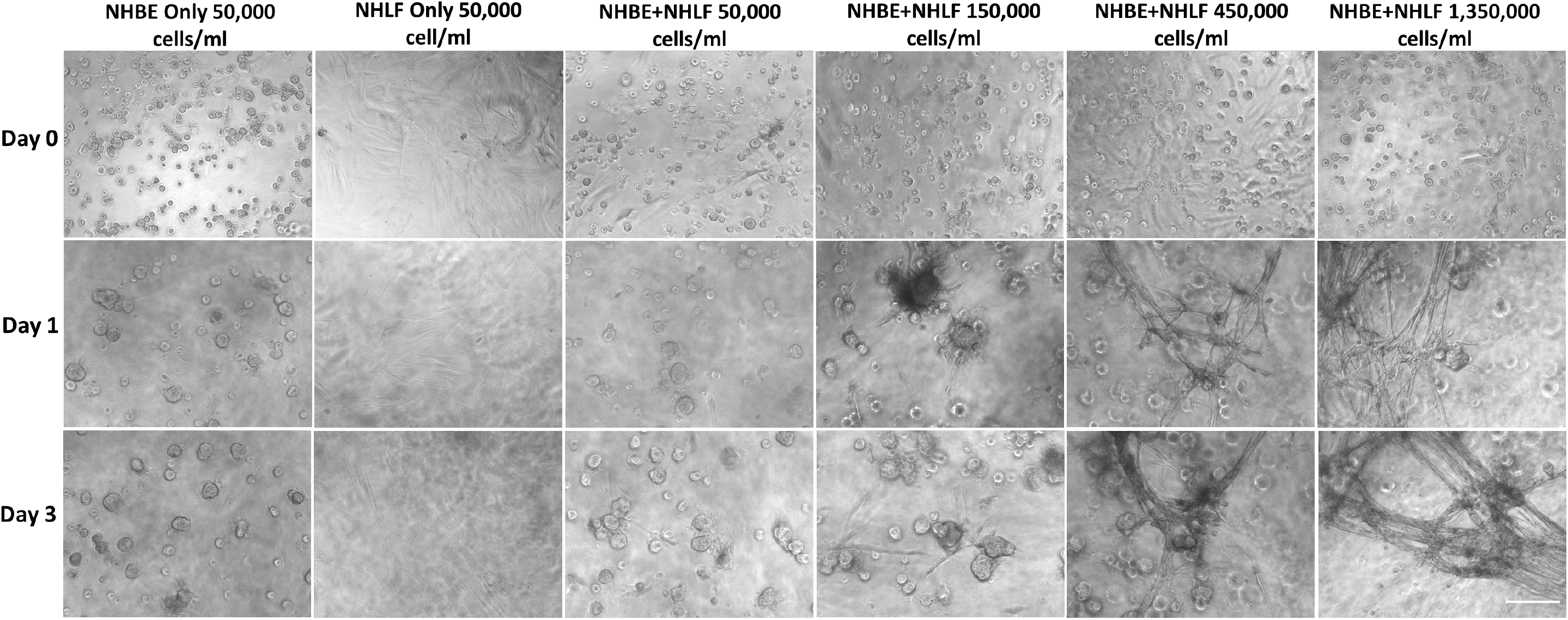
Normal healthy human bronchial epithelial (NHBE) cell and normal healthy human lung fibroblast (NHLF) co-culture: NHLF seeding density was increased from 6,000-1,350,000 cells/ml and NHLF cells allowed to adhere to the bottom of wells of a 96 well plate before being layered with 25% matrigel and then seeding NHBE cells in 5% matrigel. Increases in NHLF resulted in increased numbers of tubular structures formed by epithelial cells. Images are representative of those from 2 wells from each experiment and n=3 biological repeats. Scalebar = 100μm.

**Figure 2.**
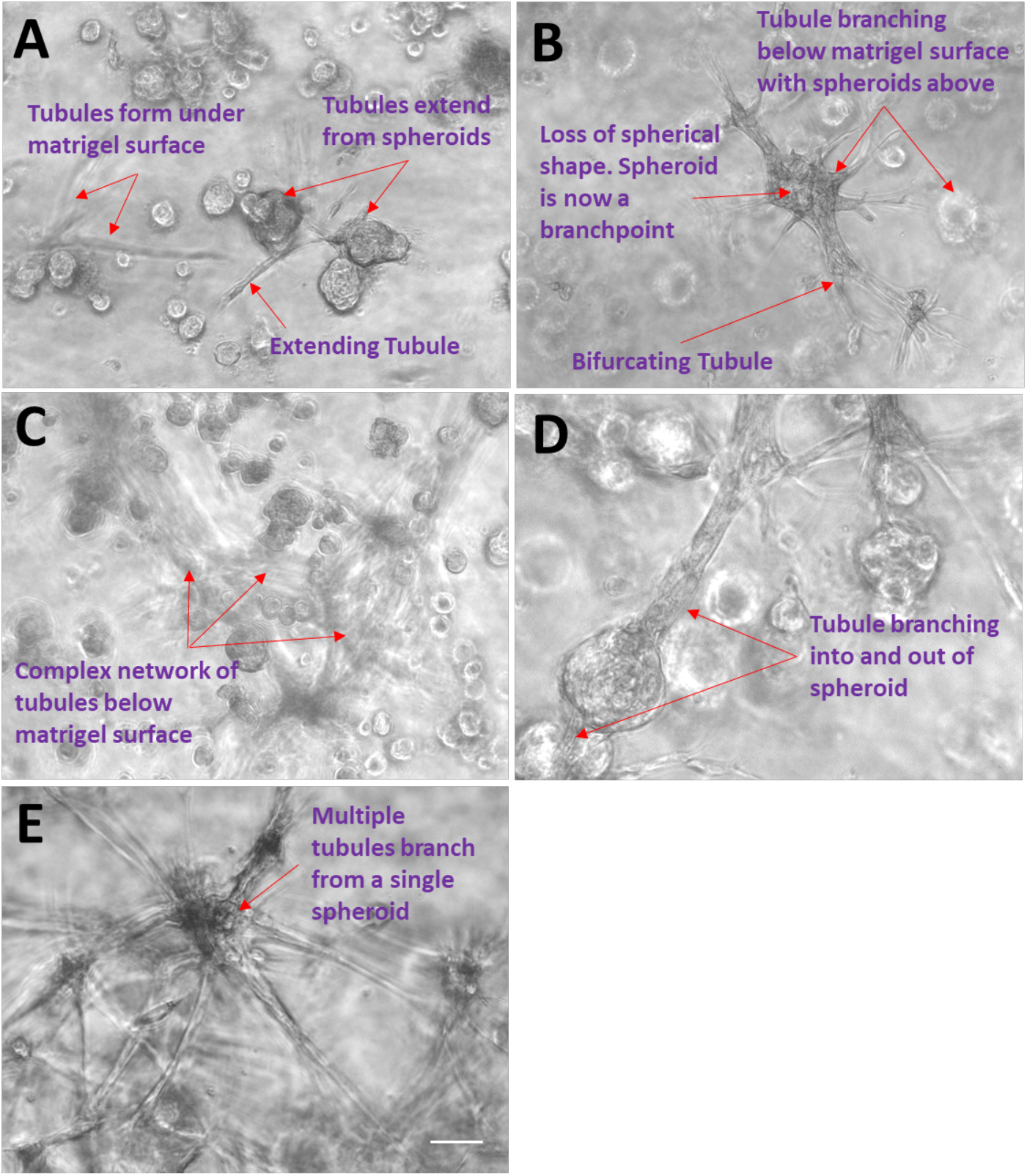
Bronchotubule formation: Normal healthy human bronchial epithelial (NHBE) cells first aggregated into spheroids (A) that grew long rod like protrusions (B) that eventually formed into an interconnected set of tubular structures that looked ganglionic (C and E). Extending tubules proximally formed a bulbous structure that branched (D). Images are representative of those seen across n=3 donors. Scalebar = 100μm.

Regular feeding was required as the media became acidic after 12hr reflecting the rapid growth of cells under these conditions. Within 3 days the rods merged together and formed lumens (**Figure 3**). We have termed these previously unreported novel structures *bronchotubules*. Further increasing the NHLF concentration to 450,000-1,350,000 cell/ml shortened the time to rod formation to 24hr (**Figure 4**). Interestingly the terminal points of the growing bronchotubules formed rounded bulb structures **(Figure 4)**. The culture survived for 4-6 days, however at this time-point, tubule structures collapsed.

**Figure 3.**
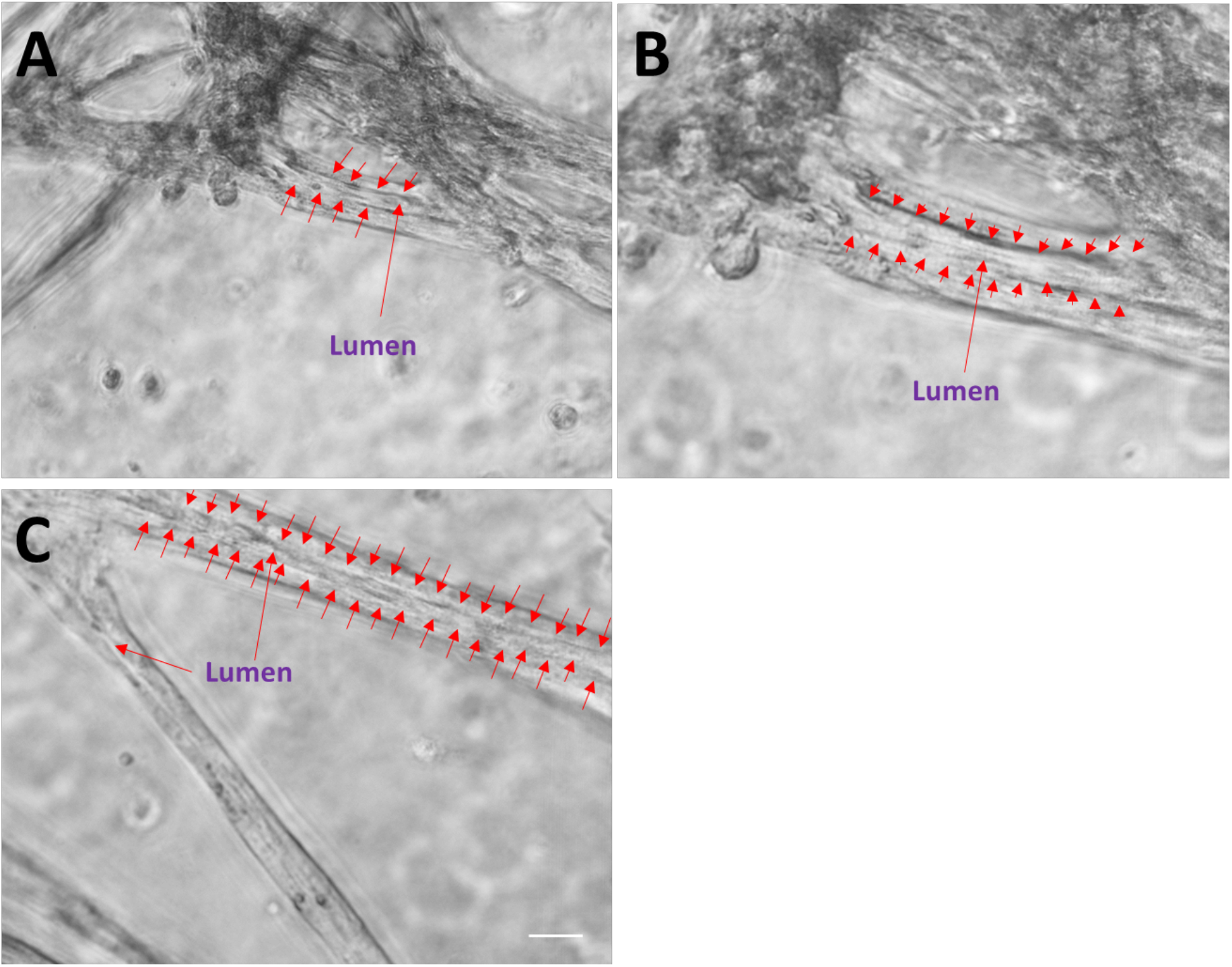
Bronchotubule morphology: Rod like structures formed lumens by Day 3 (red arrows A-C). Results are representative of bronchotubule formation in 2 wells per plate and in n=3 biological repeats. Scalebar = 100μm.

**Figure 4.**
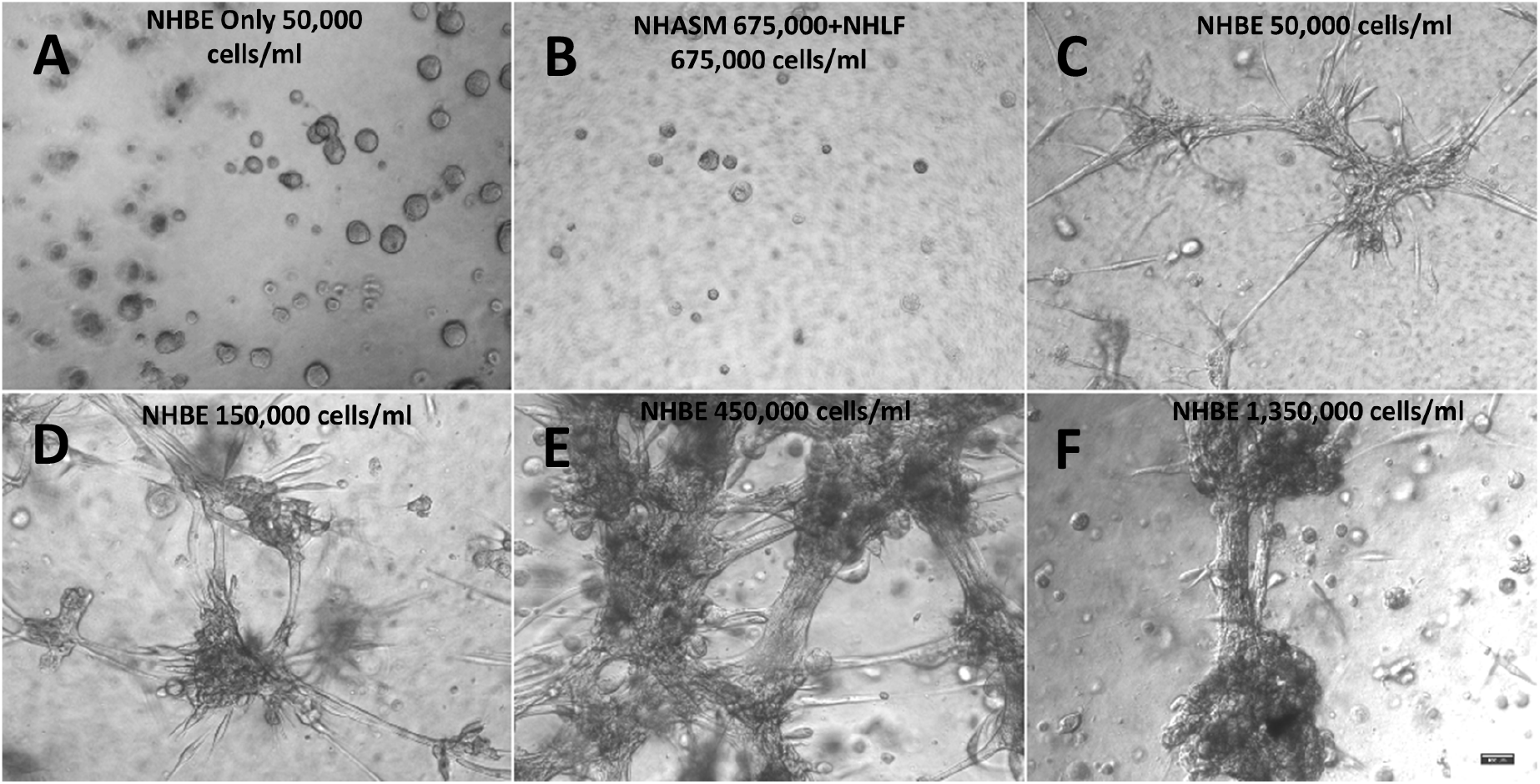
Epithelial cell (NHBE) concentration optimisation: NHBE concentration was varied while fibroblast (NHLF) and smooth muscle (NHASM) seeding density was kept constant at 675,000 cells/ml (A-F). 10x magnification, scalebar = 100μm. Images are representative of those seen across n=3 donors.

### Normal Human smooth muscle cells (NHASM) incorporate realistically into bronchotubules but do not rescue tubule collapse

Bronchiole physiology contains airway smooth muscle that bands around the tube providing structural support and control luminal dimensions. To model this micro-architecture NHASM cells were added to the culture to stabilise the tubules. The stromal cell concentrations were kept constant at 1,350,000 cell/ml (675,000 cell/ml) each for NHLF and NHASM. In control experiments in the absence of epithelial cells, stromal cultures did not branch (**Figure 4B**). However, in the triple culture, bronchotubules were once again observed and wells with the highest NHBE levels resulted in the thickest tubes (200-500μm)(**Figures 4 & 5A**).

**Figure 5.**
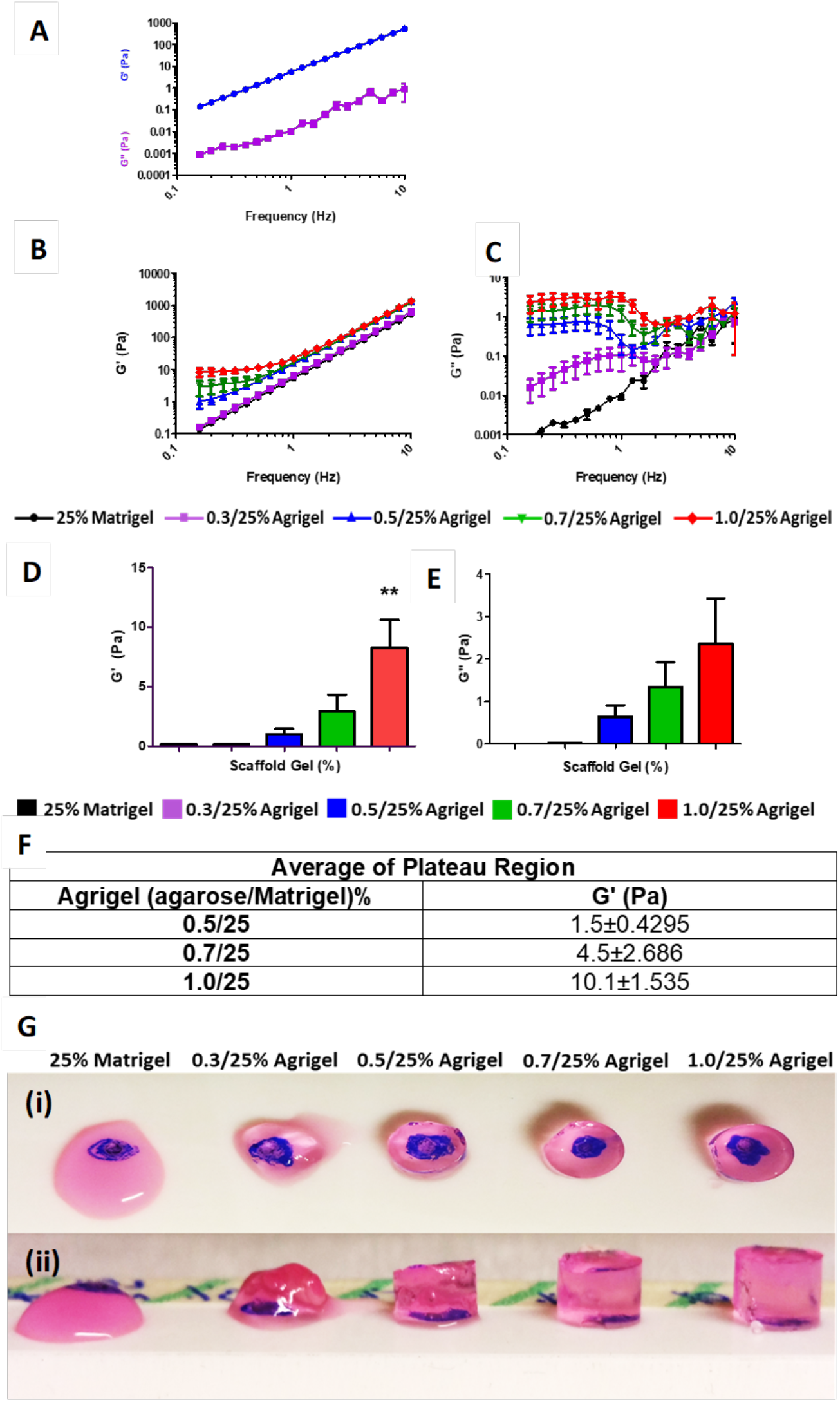
Viscoelastic properties of agrigel scaffold: (A)Storage modulus (G’) and loss modulus G” of 25% matrigel alone (B) Storage modulus and (C) loss modulus of agrigel at different concentrations(D). Storage modulus (G’) and (E) loss modulus (G”) of agrigel at different concentrations measured at 0.16Hz. Results are presented as mean±SEM of n=3 independent experiments. (F) Average G’ of plateau region showing increasing scaffold stiffness with increasing agarose concentration. (G (i)) Clarity of agrigel scaffold blue dot below gel can be clearly visualised. (G (ii)) Stiffness of agrigel increases with increasing agarose concentration.

To demonstrate the incorporation of smooth muscle functionality into the bronchotubules, we stimulated the bronchotubules with the muscarinic receptor antagonist carbochol (10^−3^M). In a pilot experiment, this caused the tubules to contract **(Figure S6B & C)**, suggesting that the NHASM are correctly incorporated into the tubules. The triple culture system incorporating smooth muscle therefore successfully recapitulates both the morphology and the contractile function of natural airway. However, the bronchotubules were not long-lived and collapsed between 4-6 days as with the simpler NHBE-NHLF co-cultures **(Figure 4)**.

### Developing a mechanically-controlled scaffold for long-term stabilization of tubules

For effective application as an organoid, it is essential to stabilize the bronchotubule structures for longer term culture, beyond the initially obtained 4-6 days. Such longer-term stability is also inherently more representative of physiological conditions. With this in mind, we hypothesize that the tubule collapse at 4-6 days was mechanically driven. Since the co-culture incorporates significant numbers of contractile cells (fibroblasts and smooth muscle) there is a natural tendency for cell-contraction to shorten and for tubules to collapse. This tendency must also exist for airway tubules in vivo and is prevented by the mechanical integrity of the surrounding ECM. Since 25% Matrigel has shown itself suitable to stimulate bronchotubule formation, we took this as a starting point, looking to modify its mechanical properties without altering the biochemistry of the material.

While the mechanical properties of biomaterials are often summarized by quoting a single number such as a Young’s Modulus the reality is more complex. In particular, biomaterials are often viscoelastic, meaning that they combine elastic or solid-like properties together with viscous or liquid-like properties. Furthermore, both viscous and elastic properties are time-dependent, such that a material can manifest quite different behaviour in response to deformation at different speeds. We have used parallel plate oscillatory shear rheometry to measure the time-dependent viscoelastic properties of scaffold candidates. This technique outputs two material properties: *the storage modulus, G’*, which stands for solid-like stiffness, and becomes proportional to Young’s Modulus for a purely solid material, and the *loss modulus, G’’*, which represents liquid-like properties and is closely related to the viscosity for a pure liquid (for a full interpretation see [30]). Both of these were determined as a function of the deformation frequency to reveal the material’s time-dependent properties.

The mechanical properties of 25% Matrigel were first measured (**Figure 5A**). It can be clearly seen that the value of the storage modulus, *G’*, is substantially greater than that of the loss modulus, *G’’*, for all frequencies measured. Hence, 25% Matrigel can be broadly characterized as a solid-like elastic material, at least within the measured range. However, the time-dependence of *G’* follows a power-law behaviour with frequency, such that at lower frequencies, its value is still decreasing with no sign of plateauing at any minimum value. Physically, this implies that 25% Matrigel shows substantial stiffness when it is rapidly deformed, however for slower, longer-term deformations, the elastic resistance to deformation falls with no lower limit, meaning that the material will be unable to resist a constant continuously applied force.

It is reasonable to divide biological forces into two (simplistically) time-scales. Firstly, individual cells are known to sense the mechanical properties of their surroundings by relatively rapid protrusion and retraction of filopodia [31–33]. Since this process is relatively rapid, 25% Matrigel could appear to cells as a significantly stiff material. However, considering large-scale structures such as the bronchotubules, the incorporation of highly contractile cell types into the structure is likely to lead to long-term constant contractile forces that operate continuously over a long period of time. In the face of such long-term forces, 25% Matrigel cannot show any resistance to very slow deformation and hence the collapse of the bronchotubules is not unexpected.

To modify the 25% Matrigel without altering the concentration of its biochemical factors, we incorporated agarose as a stiffening reagent. Agarose is a good candidate since it is biocompatible but not known to bind cell surface receptors, so that its influence on the tissue development should be purely mechanical. Furthermore, agarose gels set solid with decreasing temperature and do not require chemical cross-linking agents in contrast to other stiffening agents such as alginate which sets after addition of calcium. The mechanical properties of agarose-25% Matrigel mixed gels, which we term *“agrigels”* were measured using parallel-plate rheometry (**Figure 5B and C**). The agrigels show, *G’* > *G’’* at all frequencies, although the gap between the storage and loss moduli narrows at low frequency where these must be considered truly viscoelastic materials. However, the low frequency behaviour of the storage modulus, *G’*, is very different from pure Matrigel (**Figure 5B**). Rather than continuing to decrease as a power law with decreasing frequency (straight line on plot), G’ deviates upwards, approaching a plateau value (**Figure 5B**). This plateau is clearly seen for 0.7% agarose and is a reasonable extrapolation for 0.5% and 0.3%. The presence of a low-frequency plateau is highly significant, implying that the material will be able to support a long-term constant force and thus could provide a viable scaffold for large-scale contractile tissues such as the bronchotubule system (**Figure 5B**). The values of stiffness (*G’*) was controlled by the quantity of added agarose: the value of *G’* in the plateau region increased by up to 10,000-fold compared with pure 25% Matrigel with agarose concentrations from 0.3–1.0% (**Figure 5D, E and F**). Intriguingly this was a much greater range of stiffnesses than was seen using pure agarose gels at the same concentrations (**Supplementary Figure S1**). This ability to resist long-term forces was qualitatively confirmed by the ability of agrigel constructs to retain their shape under gravity better than pure 25% Matrigel (**Figure 6G**).

**Figure 6.**
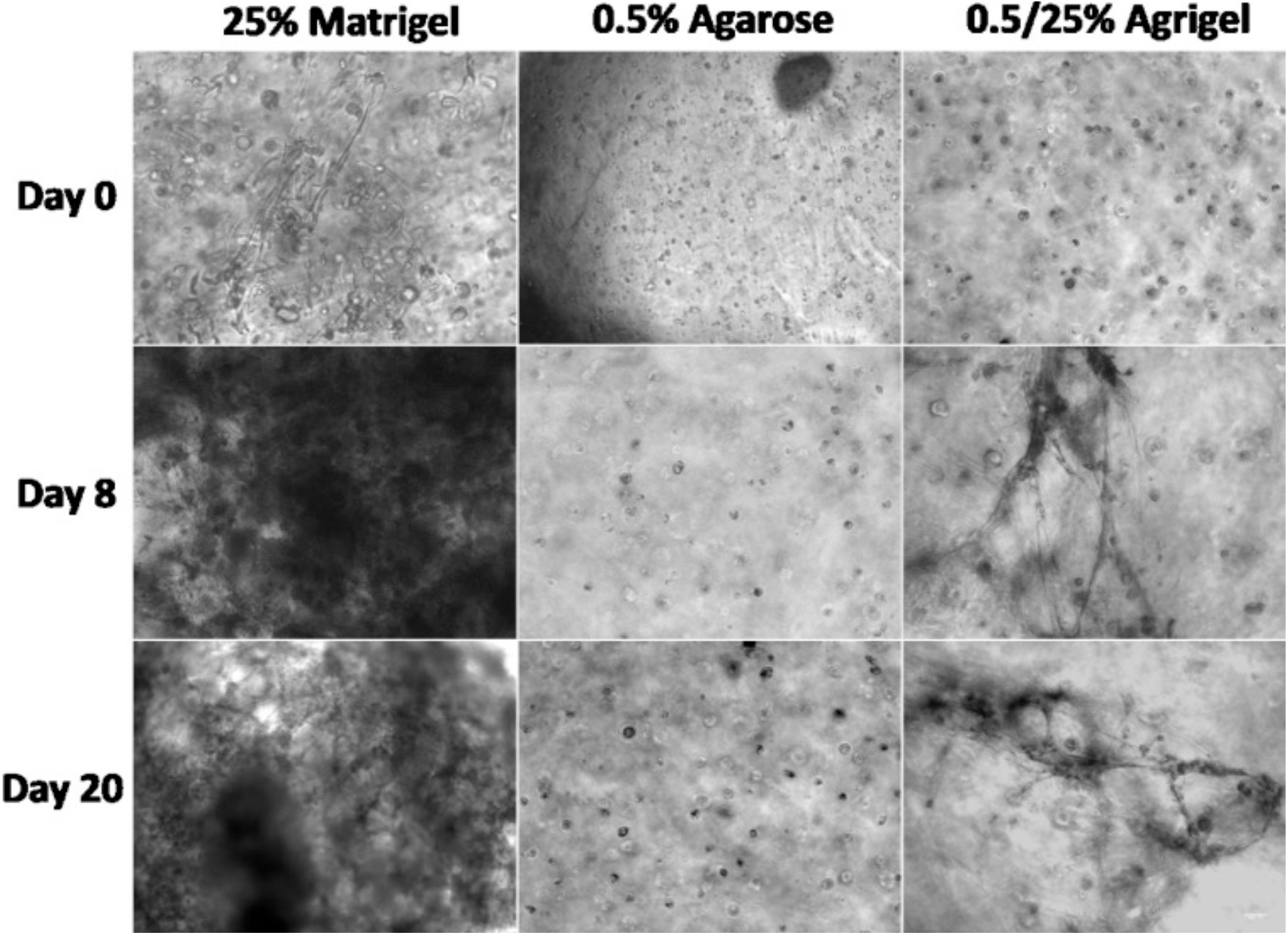
Scaffold optimisation: Triple culture of normal healthy human bronchial epithelial cells (NHBE), normal healthy human lung fibroblasts (NHLF) and normal healthy human airway smooth muscle (NHASM) cells in different scaffolds. Cell seeding density; NHBE = 450,000 cells/ml, NHLF = 675,000 cells/ml and NHASM = 675,000 cells/ml. 10x magnification, scale bar = 100μm. Images are representative of n=3 using cells from the same patient.

We thus consider the agrigel a well-suited novel scaffold system for 3D culture of complex-structured contractile organoids, demonstrating the key ability to support low-frequency/long-time forces and with stiffness control across 4 orders of magnitude without changing the underpinning biochemistry of the matrix.

### The increased stiffness of the agrigel scaffold enables long term culture of bronchotubules without collapse

To assess whether matrix stiffness affected the durability of organoid cultures we examined whether addition of agarose to enhance matrigel viscosity had an impact on bronchotubule formation and sustainability. Bronchotubules were grown in 0.5 and 0.7% agrigel. The tubules observed in 0.7% agarose containing agrigel looked thinner compared to that of 0.5% agrigel **(Figure 6)** and were unable to form a complex network, with spheroidal structures the predominant morphology.

Based on these results and that of the rheological analysis **(Supplemental Figures S2 & S3),** we tested agrigel containing 0.5% agarose (0.5%-agrigel) to provide an appropriate balance between the stiffness necessary to support the long-term contractile forces while also delivering a sufficiently compliant gel to allow for cell-migration and tubule growth. In 0.5%-agrigel, bronchotubules successfully formed after 8 days, and survived until the end of the experiment at 20 days. Bronchotubules grown in 0.5% agrigel resembled morphologically the airway *in* vivo. In contrast, cells grown in 0.5% agarose alone were unable to form tubular or spheroidal structures indicating the critical role of the Matrigel **(Figure 7)**. This data shows that the mechanical properties of the matrix influence tubule formation with the bifurcating tubules growing unidirectionally rather than undertaking non-specific branching in all directions. All further experiments were performed using 0.5%-agrigel.

**Figure 7.**
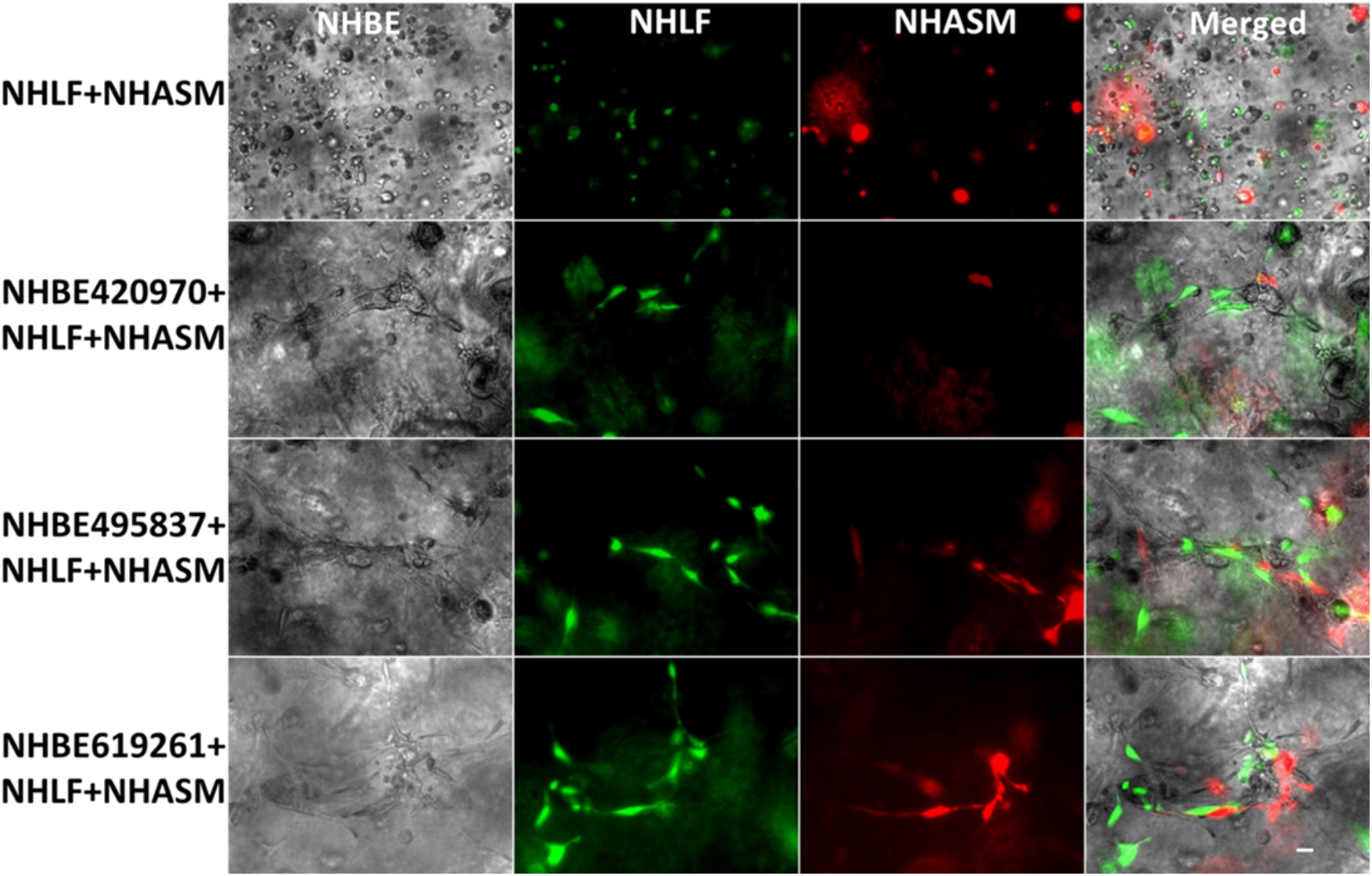
Organization of culture-generated tissue: Bronchotubule Triple Culture in 0.5/25% Agrigel: Triple culture of lentivirally transduced of normal healthy human lung fibroblasts (NHLF, green) and normal healthy human airway smooth muscle (NHASM, red) cells at 675,000 cells/ml respectively with normal healthy human bronchial epithelial cells (NHBE) at 450,000 cells/ml. NHLF and NHASM do not form tubules in agrigel compared to triple culture which form tubules over 20 days cultures. Magnification 10x, Scale bar is 100μm. Images are representative of those from n=3 different donors.

Next, we determined whether culture with different cell types formed the correct airway architecture. We expected that NHBE would be surrounded by NHLF and that NHASM cells would band across the outer edge of the bronchotubule. To elucidate the position of NHLF and NHASM in bronchotubule structure in 0.5/25% agrigel, NHLF and NHASM were labelled with lentiviral-transfected yellow fluorescence protein (YFP) and mCherry respectively **(Supplemental Figure S4)**. The culture was repeated with primary NHBE from 3 healthy donors. NHBE cells (grey) were surrounded by NHLF (green) with NHASM cells (red) forming the outer edge of the tubules mimicking the human airway and indicating that epithelial-stromal stratification occurs in this model **(Figure 7–8)**. Tubules from each NHBE cell donor were maintained for 20 days although cells from each patient formed morphologically different bifurcating fractal structures where bronchotubules from each patient generated daughter branches of different size, length and shape **(Figure 8)**. Epithelial cells that did not migrate into the gel and interact with other stomal cells formed spheroids but not bronchotubules **(Supplemental Figure S5)**.

**Figure 8.**
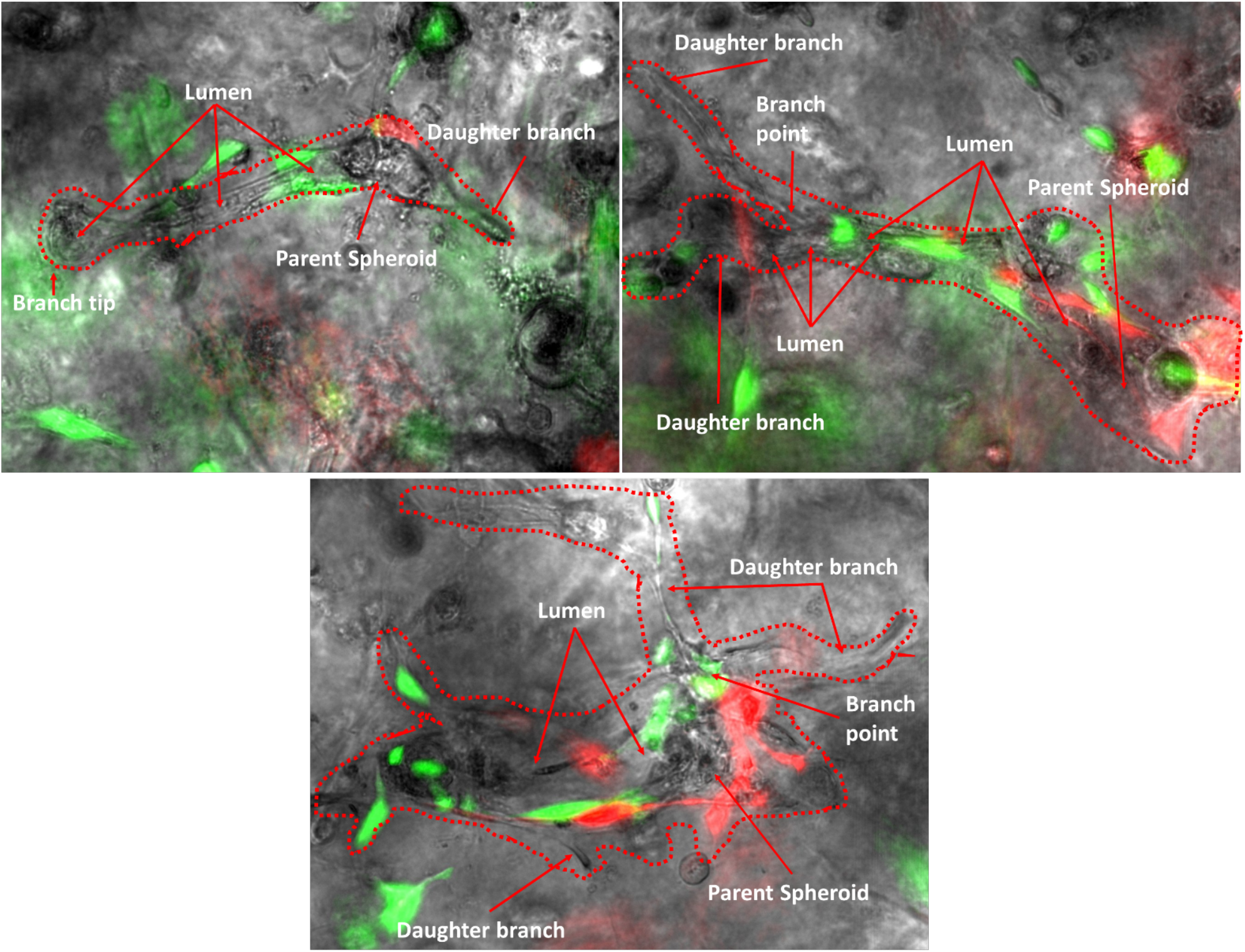
Organization of culture-generated tissue: Bronchotubule Triple Culture in 0.5/25% Agrigel Day 20: Bronchotubules arise from spheroids and form bifurcating branches over 20 days culture. Normal healthy human lung fibroblasts (NHLF, green) and normal healthy human airway smooth muscle (HASM, red) cells surround and support the growth of normal healthy human bronchial epithelial cells (NHBEC, grey). Images are representative of n=3 experiments using NHBECs from 3 different donors. Scalebar = 100μm.

These results show that bronchotubule culture is able to recapitulate the airway cellular architecture and that distinct cell lineages are autonomously able to migrate and organise themselves into expected regions within the developing bronchotubule even though they originate from 3 separate healthy donors. The presence of a cellular lumen indicates that epithelial cells in the inner lining of the tubules are undergoing apical-basal polarisation observed in luminogenesis. However, bronchotubule formation is not only cell-dependent, but requires an ECM with the right level of viscoelasticity. We have shown that ECM stiffness is crucial in patterning and holding the structure **(Figure 7 and 8)** which collapsed after 4 days in Matrigel alone. These experiments also suggest that primary human airway cells are inherently programmed to form the correct airway patterning when exposed to the right ECM mechanical niche.

## Discussion

We report here for the first time, the generation of functional contracting lung tubular branching NHBEC-derived organoids in the presence of stromal NHLF and NHASM cells, that we have termed bronchotubules. We introduced a new scaffold material, agrigel, whose time-dependent mechanical properties were designed to allow long-term culture of these contractile organoids. To our knowledge, agarose in combination with Matrigel has not previously been used as a mixed gel scaffold for organoid culture, although applications of collagen-agarose mixed gels are reported [34–36]. Bronchotubule organoids developed in this study, display epithelial-stromal stratification and cannot form without epithelial-stromal signalling.

Previous studies have reported that mixed stromal-epithelial cell populations can form spheroidal structures [17] that branch and lose their spheroidal identity. Tubules formed in 25% matrigel merge into each other as seen in our bronchosphere culture after 4 days whereas tubules cultured in the stiffer 0.5%-agrigel continue branching in a unidirectional manner with adjacent tubules not merging into each other. Bronchotubules grown in 0.5% agrigel do not appear to reach quiescence and continue to branch and grow for up to 3 weeks – the maximal duration of analysis.

During lung development *in vivo*, branching ceases when tip cells begin to form alveolar structures with epithelial cells acquiring a SOX9/ID2+ identity and eventually forming ATI and ATII epithelial cells [9, 37, 38]. This SOX9/ID2+ phenotype is theorised to arise from tip cells of nascent bifurcating tubules. Our study did not include alveolar populations and perhaps the addition of ATII cell populations are required to give a stop signal to branching and enable the formation of alveolar structures [4, 9, 20, 38]. The formation of tubules may indicate that epithelial cells are pre-programmed to create tubular structures in certain microenvironments.

Organoid culture methods have applied various matrix scaffolds such as matrigel or collagen gels [17, 20, 39, 40]. Furthermore, the concentration of scaffold gel used varies in literature which may have major effects on the resultant organoids formed. We have shown that even a 0.2% change in agarose concentration has a profound effect on the type of structures that form. Varying this quantity may unwittingly increase or decrease exogenous proteins that stimulate organoid assembly. Maybe the level of resistance of very stiff 0.7% gel to cellular migration was too high resulting in less tubular and more spheroidal structures. Studies where epithelial cells grown in 50% matrigel results in tracheosphere organoids that are clonal arising from the division of a single progenitor cell [41] contrary to 25% matrigel formulations where organoids develop by the migration and aggregation of cells. The role of cellular migration and rearrangements *in vivo* are crucial for lung branching morphogenesis during development. For example, [42] have shown that stromal progenitor migration to the tips of nascent lung tubes and their differentiation into smooth muscle cells is crucial for of tubular bifurcation. This requires the surrounding matrix to be compliant and viscous enough to allow cells to pass indicating that the matrix plays a major role in cellular migration and that there is an optimal fitness for cellular ability to migrate through matrix [43]. Whilst comparing organoid behaviour across studies is difficult it is evident that the matrix scaffold used is important.

This study has several limitations including the sole use of structural cells from healthy subjects and exclusion of immune cells. Culturing bronchotubules using diseased lung cells such as COPD-HBEC may generate aberrantly branching tubules due to dysregulated cell-cell signalling. If cells are preprogramed to make these structures, diseased lineages should recapitulate abnormal development. Furthermore, disease-healthy cell hybrid bronchotubules such as NHBE with ASMs and COPD fibroblasts or COPD HBECs with NHLF and NHASM cells could help elucidate disease driving cell lineages of abnormal developmental processes and whether one could rescue bronchotubule phenotype by replacing the disease driving lineage with a healthy lineage. Furthermore a recent study showed that healthy cells grown in matrix from diseased patients result in organoids that recapitulate diseased lung architecture [44]. Future studies should examine the differentiation of epithelial cell subtypes in bronchotubule cultures over time using single cell RNA-sequencing [45] comparing the data with bronchial biopsies from healthy and COPD donors.

In summary, this method has enabled long-term stable culture of the present bronchotubule system. Since the mechanical microenvironment can easily be varied by specifically increasing or decreasing the agarose concentration, it will be possible to mimic different scaffold environments of different airways diseases like COPD and IPF. The bronchotubule system opens the possibility to investigate questions around the role of cell-matrix interaction and cell-cell communication in tissue patterning, repair and disease *in vitro*. We anticipate the use of hybrid cultures to investigate the interactions between diseased and healthy cells. Further advancement of organoid models like these will expand our knowledge of how cell-cell or cell-matrix interactions drive the development and repair of lung tissue.

## Methods

### Agrigel Formulation

Matrigel and agarose were mixed in differentiation media (1:1 Dulbecco’s Modified Eagle’s Medium (DMEM, Sigma, Poole, UK) and Bronchial Epithelial Basal Medium (BEBM, Lonza, Slough, UK) with growth factors from the Lonza bullet kit excluding retinoic acid (RA) and triidothyrine) supplemented with 100nM RA) to a final concentration of 0.5% or 0.7% agarose and 25% matrigel to form Agrigel (Agarose/Matrigel). Matrigel was stored on ice until used and was mixed with differentiation media before addition of agarose. Pipette tips were warmed and cut prior to mixing of agarose and matrigel. Agarose was used at 45°C. Gels were mixed as quickly until pink as matrigel is solid at 10°C whilst low melting agarose is solid at 36°C.

### Cell Culture

Never-smoker human airway epithelial cells and culture media from Lonza (Basel, Switzerland) were cultured according to company’s instructions and used at passage 2 (P2). Normal Human Lung Fibroblasts from Promocell (Heidelberg, Germany) were cultured in accordance with the company’s instructions and were used at P4-P5. Normal Human Airway Smooth Muscle cells donated by Dr. Charis Charalambous (NHLI, Imperial College) were cultured in Dulbecco’s Modified Eagles Medium (DMEM, Sigma) supplemented with 5% L-glutamine (200nM, Sigma) and penicillin/streptomycin (0.1 mg/ml, Sigma) and used at P4-P5 as described [46].

#### NHBE-NHLF Cell Co-culture

NHLF were seeded onto 96 well plates at either 25,000, 75,000, 225,000 or 675,000 cells/ml in 0.1ml differentiation media and adhered overnight. Media was removed, and cells were coated with 25% matrigel in differentiation media containing 100nM RA for 2h at 37°C. NHBE cells were seeded at 50,000 cells/ml in 5% matrigel in differentiation media supplemented with 100nM RA and fed every 2 days.

#### NHBE-NHLF-NHASM Cell Triple culture

NHBE, NHLF and NHASM cells were trypsinised and centrifuged at 220 x g and were mixed in 50ml falcon tubes: NHBE 45,000 cells/well and NHLF and NHASM at 67,500 cells/well. Agrigel solution was mixed with cells and pipetted into transwells precoated with 0.5/25% Agrigel. Differentiation medium with 100nM RA was added to basal wells. The culture was incubated at 37°C for 20 days. Media was replenished every 2 days.

### Lentiviral Transduction

Yellow fluorescence protein (pLEX_970_puro_DEST_YFP gifted by William Hahn (Addgene plasmid # 45295)) and mCherry (plv_mCherry gifted by Pantelis Tsoulfas, (Addgene plasmid # 36084)) plasmid in transfected bacteria were amplified and extracted with Qiagen Maxiprep kit (Qiagen, Manchester, UK). HEK293FT cells (gifted by Dr. Carlos Lopez Garcia Crick Institute) were transfected with YFP or mCherry, psPax2 and pMD.2G viral RNA plasmid envelope in OPTI-MEM with lipofectamine 2000. Lenti-X™ qRT-PCR Titration Kit (Takara biosciences) was used to calculate the viral multiplicity of infection (MOI) of 3 that was used to infect NHLF and NHASM cells.

### Microscopy

Organoids were visualised with Leica DMI6000 microscope and analysed with ImageJ version 1.52e (https://imagej.net/).

### Mechanical Characterization of Gels

50μl of either 25% matrigel, Agarose (0.3, 0.5, 0.7 or 1.0%) or Agrigel (0.3/25, 0.5/25, 0.7/25 or 1.0/25%) solution was instantly poured into a Teflon mould that was 8mm diameter and 5mm in height. Gels were solidified at 37°C/5% CO_2_ for 20 minutes and incubated for 24h in differentiation media with all supplements. At the time of measurement Rheometer discovery HR1 (TA Instrument, using TRIOS software, New Castle, DE, USA). base plate was heated to 37°C, gels were removed and compressed with a 8 mm diameter compressive plate. Storage modulus (G’) and loss modulus (G”) were acquired from oscillatory shear measurements (frequency 0.1-25rad/s).

### Statistics

One-way ANOVA with a Bonferroni’s multiple comparison test was used to assess statistically significant differences between means of two groups. All statistical analyses were performed in GraphPad Prism 6.0 software.

## Acknowledgements

We acknowledge funding from the MRC Industrial CASE Award Scheme (MR/L015854/1), the Wellcome Trust (093080/Z/10/Z), the Dunhill Medical Trust (R368/0714) and EPSRC (EP/M000044/1).

## Supplementary Figures

**Supplementary Figure S1.**
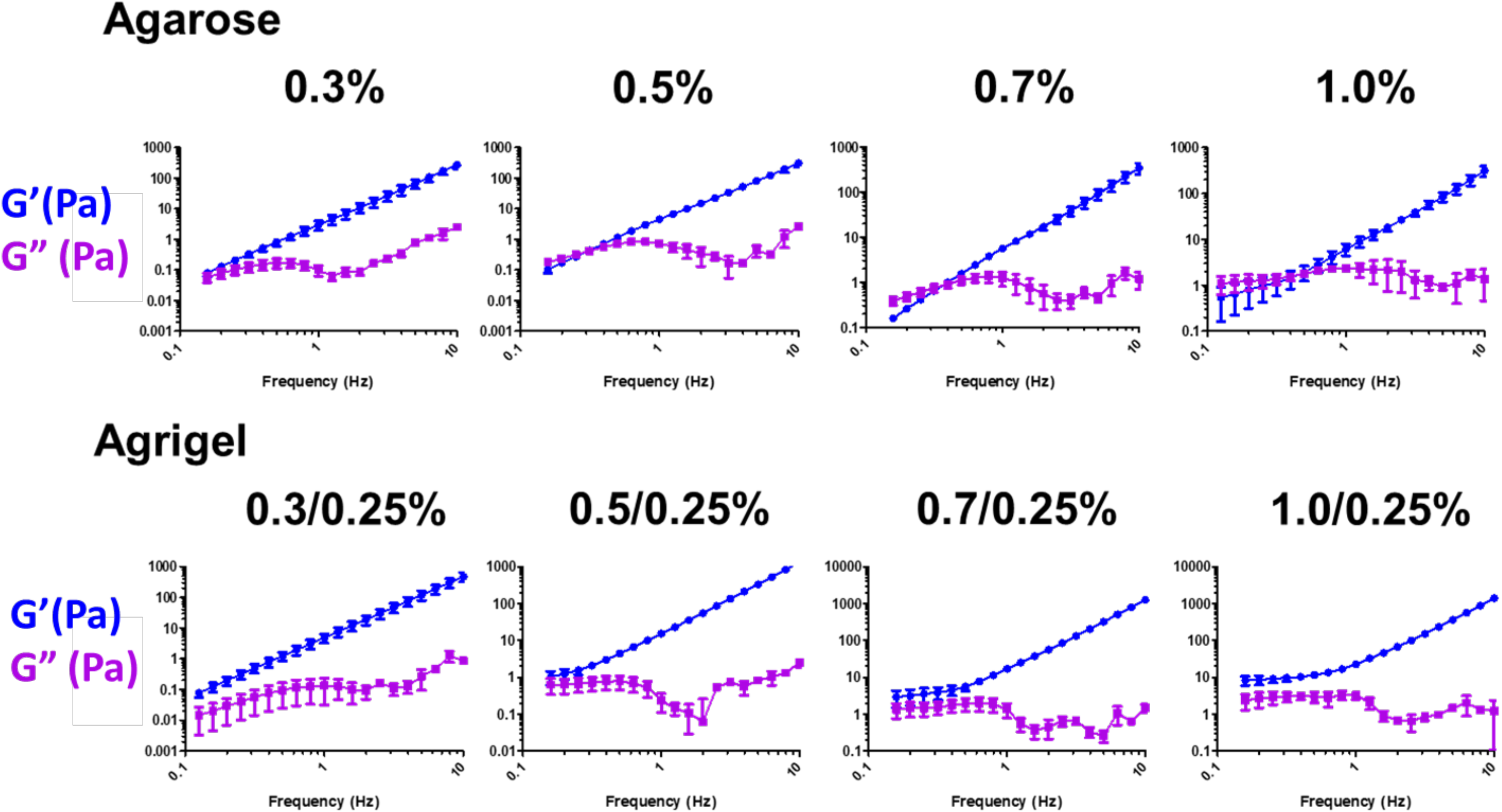
Storage modulus (G’) and loss modulus G” of different agarose or agrigel (agarose/matrigel) mixtures. Results are presented as mean±SEM of n=3 independent experiments.

**Supplementary Figure S2.**
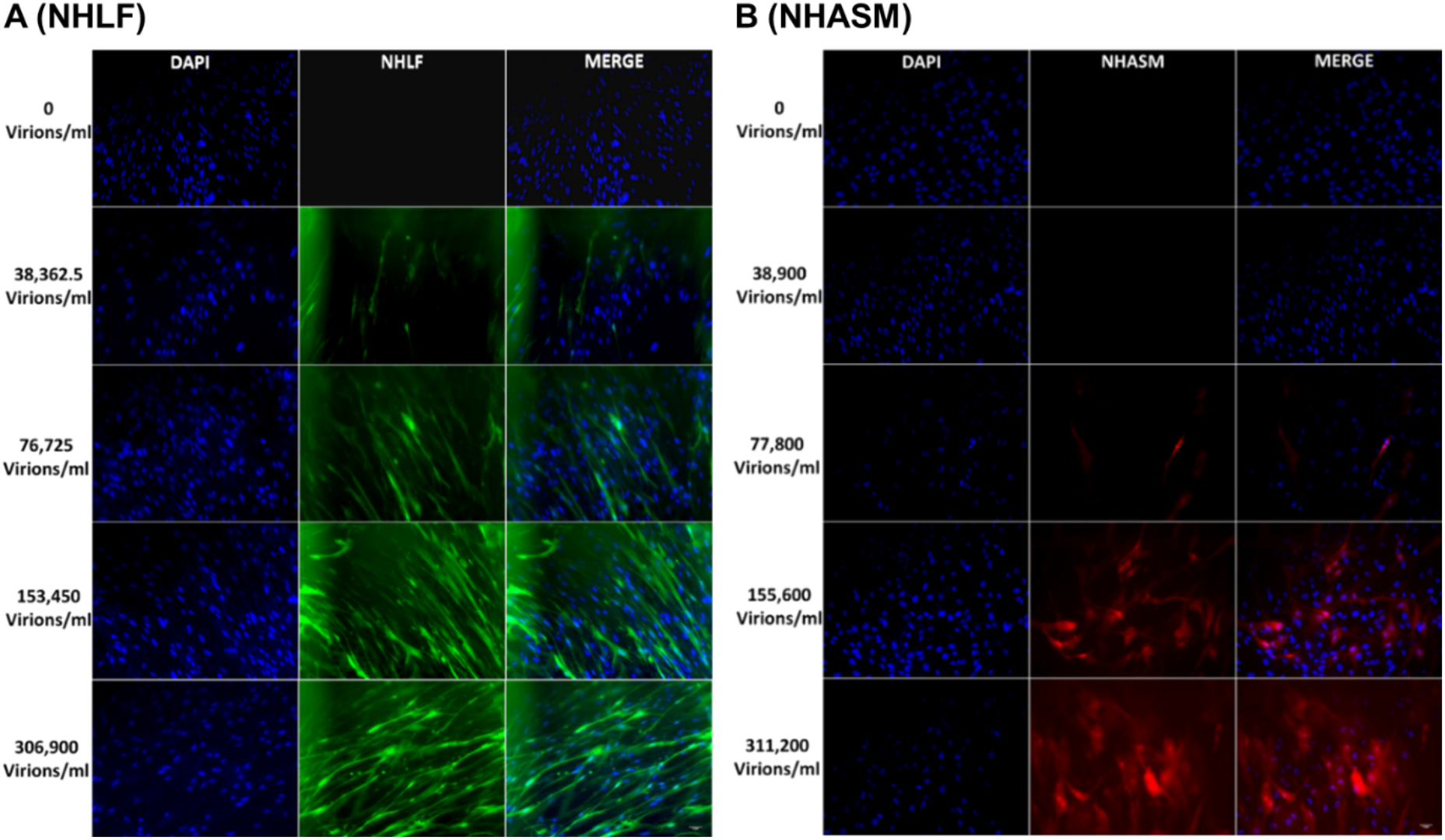
Stromal Cell Lentiviral Transduction: 100, 000 normal healthy human lung fibroblasts (NHLF) and normal healthy human airway smooth muscle (NHASM) cells were transfected with 306,900 YFP and 311,200 mCherry viral particles respectively in 1ml media with a multiplicity of infection (MOI) = 3. Images are at 5x magnification, Scale bar is 50μm. Images are representative of those from n=3 independent experiments.

**Supplementary Figure S3.**
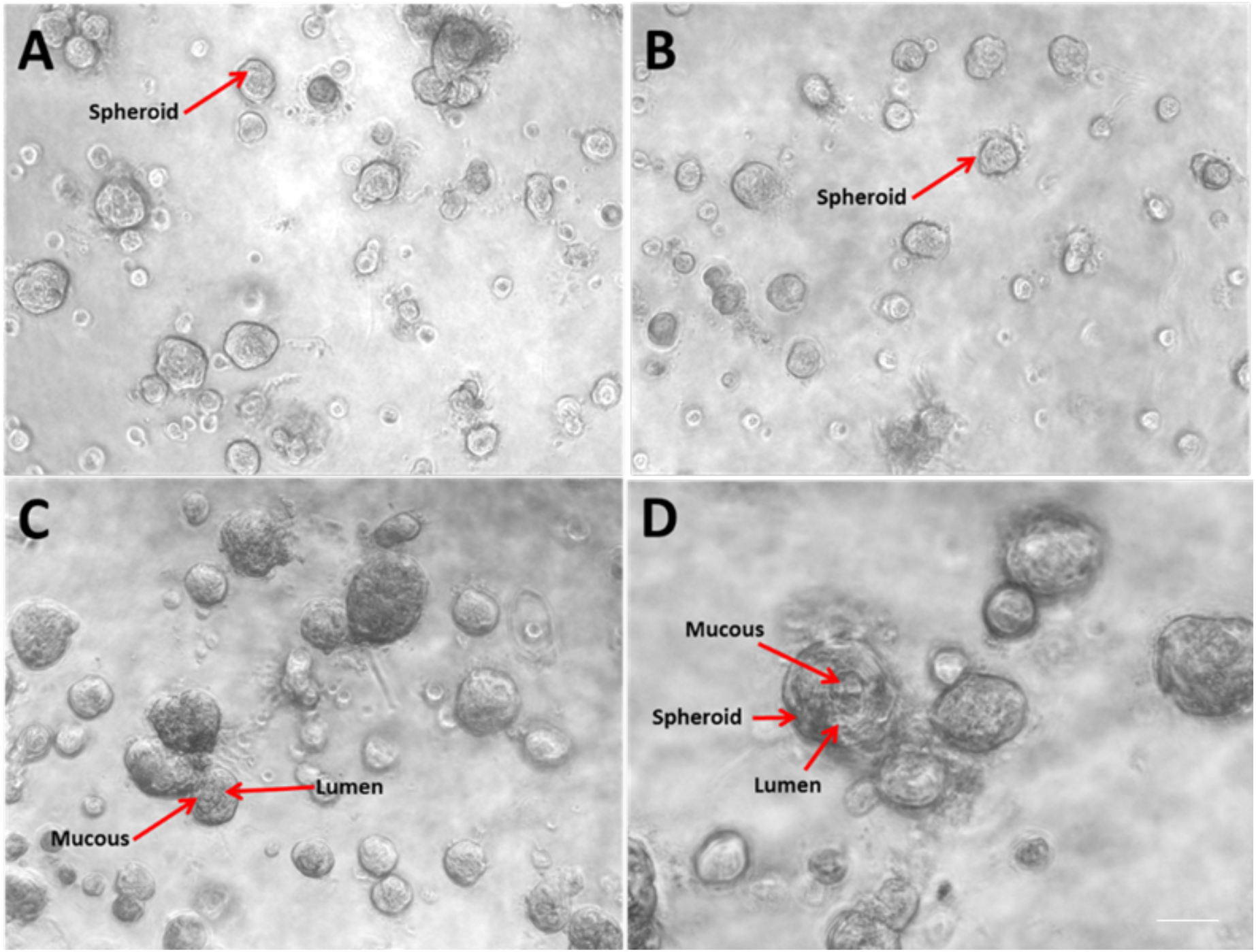
Epithelial cell behaviour above gel: Epithelial cells that did not migrate into the gel formed into spheroids and did not form rod-like or tubular structures. Images are representative of those seen in 2 wells per plate and for n=3 donors. Scalebar = 100μm.

**Figure S4.**
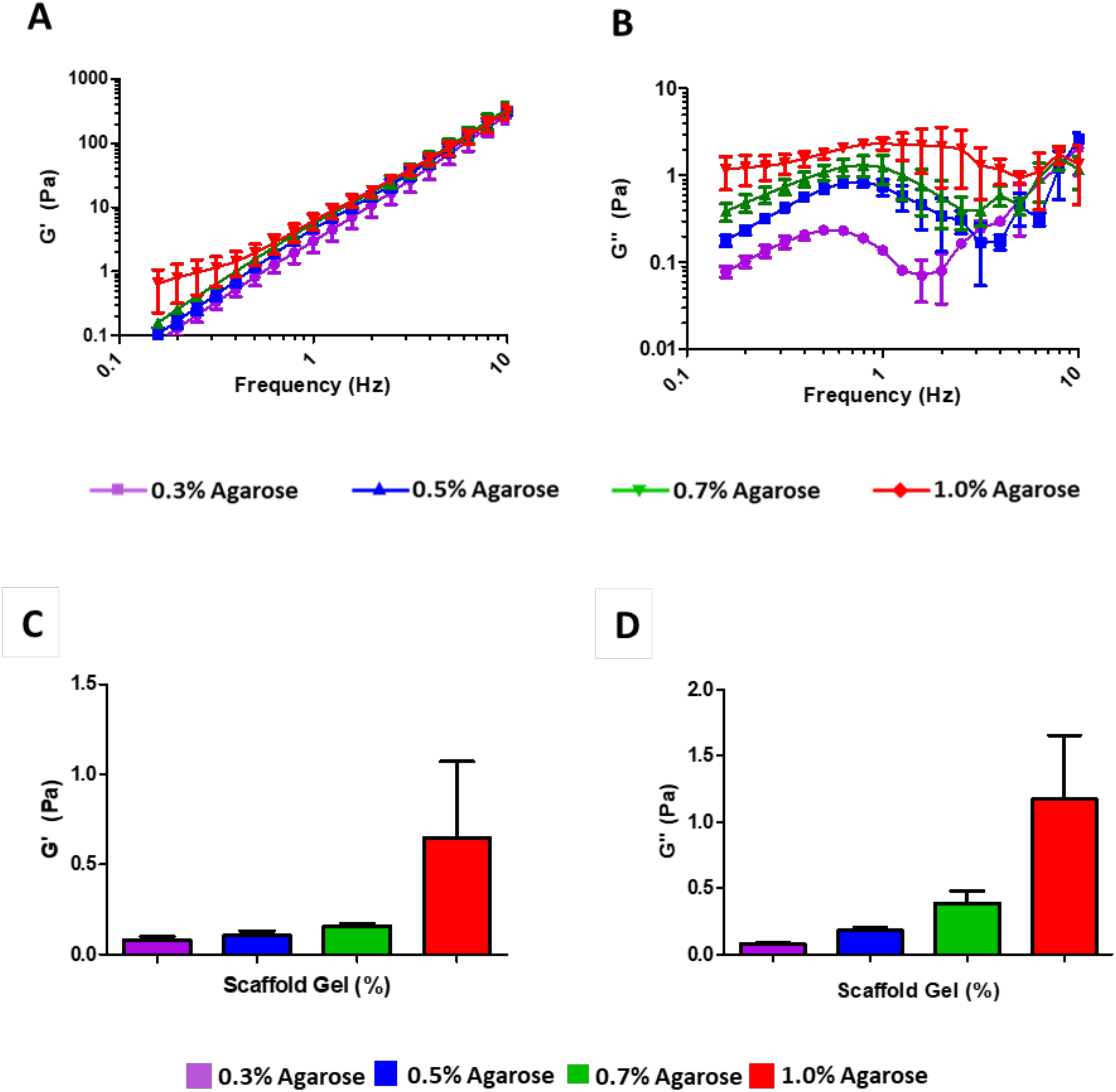
Viscoelastic properties of agarose gel scaffold: (A) Storage modulus (G’) and loss modulus G” (B) of agarose at different concentrations. (C) Storage modulus (G’) and (D). loss modulus (G”) of agarose at different concentrations measured at 0.16Hz. Results are presented as mean±SEM of n=3 independent experiments.

**Figure S5.**
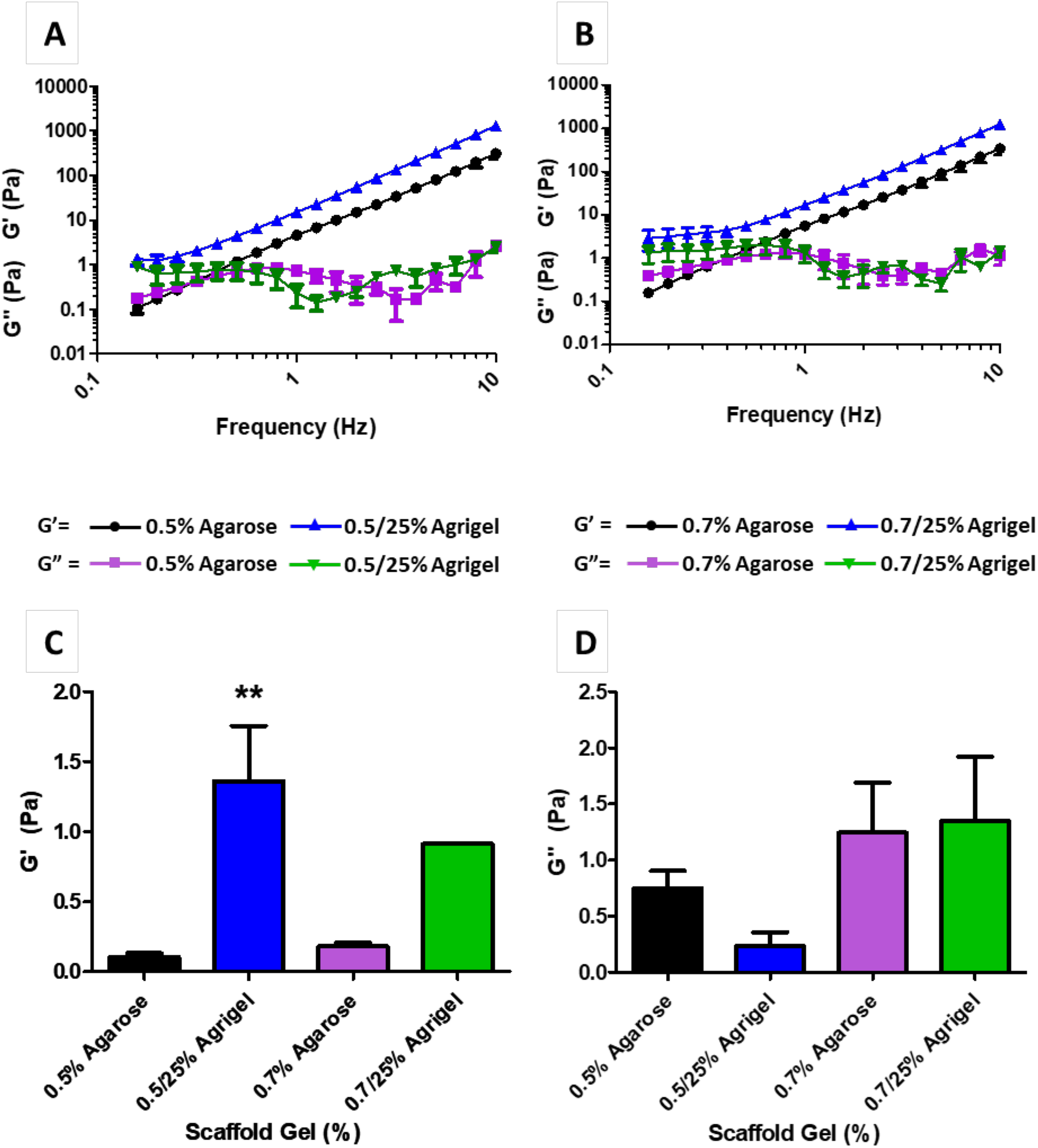
Comparison of viscoelastic characteristics of agarose and agrigel: (A) Storage and loss modulus of 0.5% agarose compared to 0.5%/25% agrigel. (B) Storage and loss modulus of 0.7% agarose compared to 0.7%/25% agrigel. (C) Storage modulus of agarose at different concentrations measured at 1Hz. (D) Loss modulus of agarose at different concentrations measured at 1Hz. All data is presented as mean SEM of n=3 independent experiments using ANOVA compared to agarose. *p<0.05, **p<0.01, ***p<0.001.

**Figure S6.**
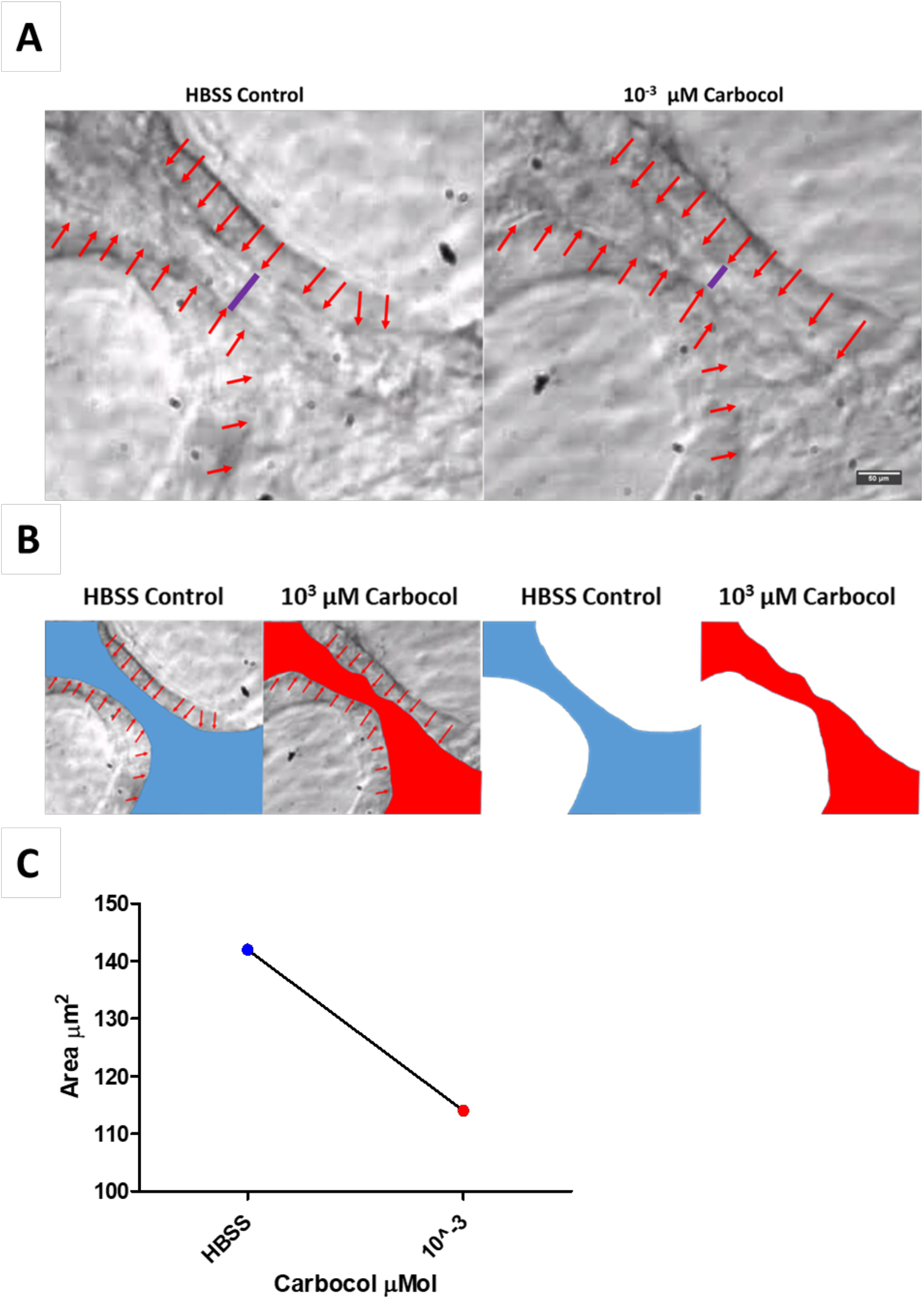
Normal healthy human airway smooth muscle (NHASM) Contraction assay: The functionality of NHASM cells was measured by the addition of the muscarinic agoninst carbachol (10^−3^M) which causes ASM contraction and airway constriction compared to Hank’s Balanced Salt Solution (HBSS) buffer control. Purple bar shows luminal contraction, A, area of lumen was measured using imageJ by marking the inside lumen (blue and red), B and C. 40x magnification, Scale bar scalebar= 100μm. n=1.

## STAR Methods

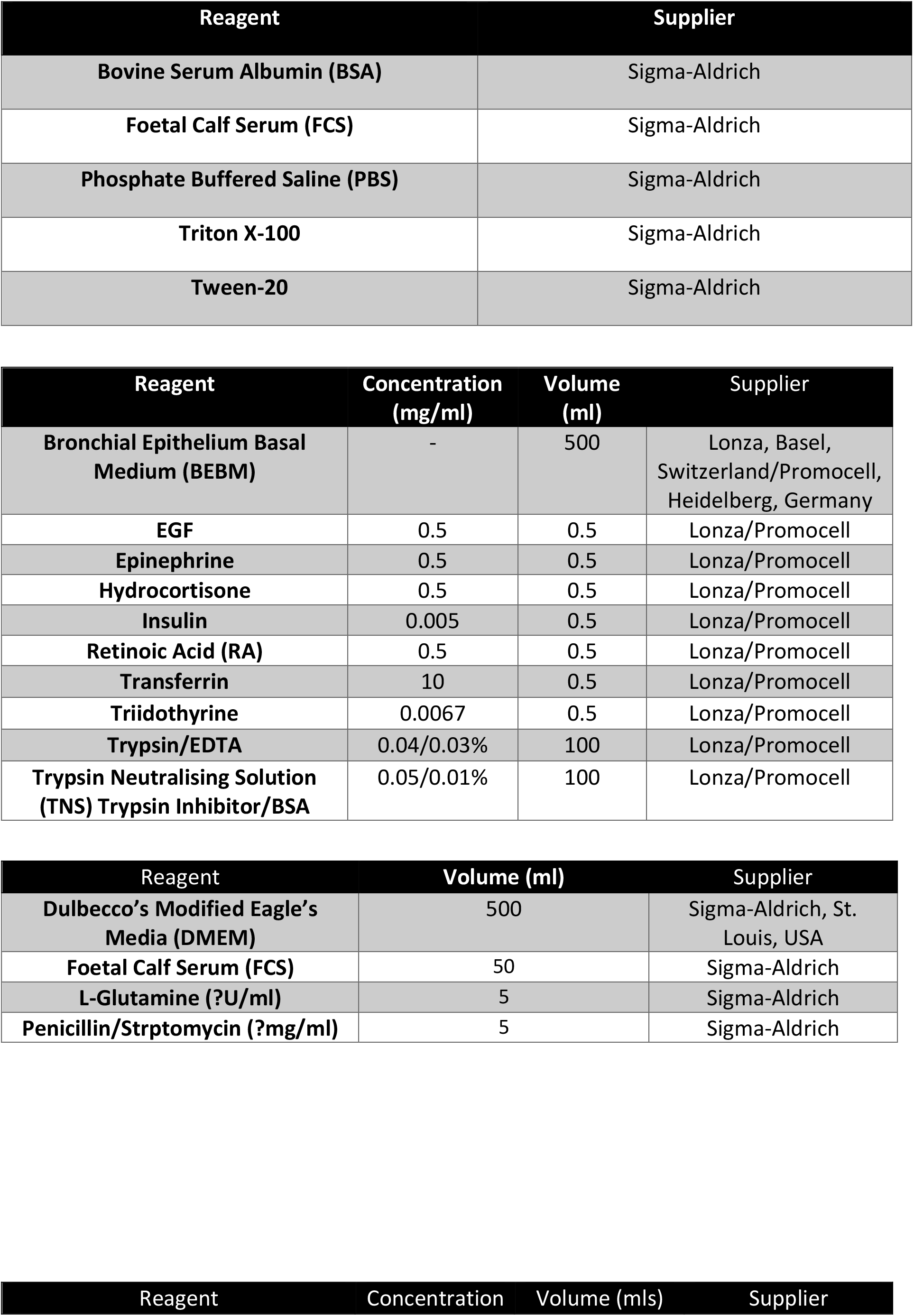
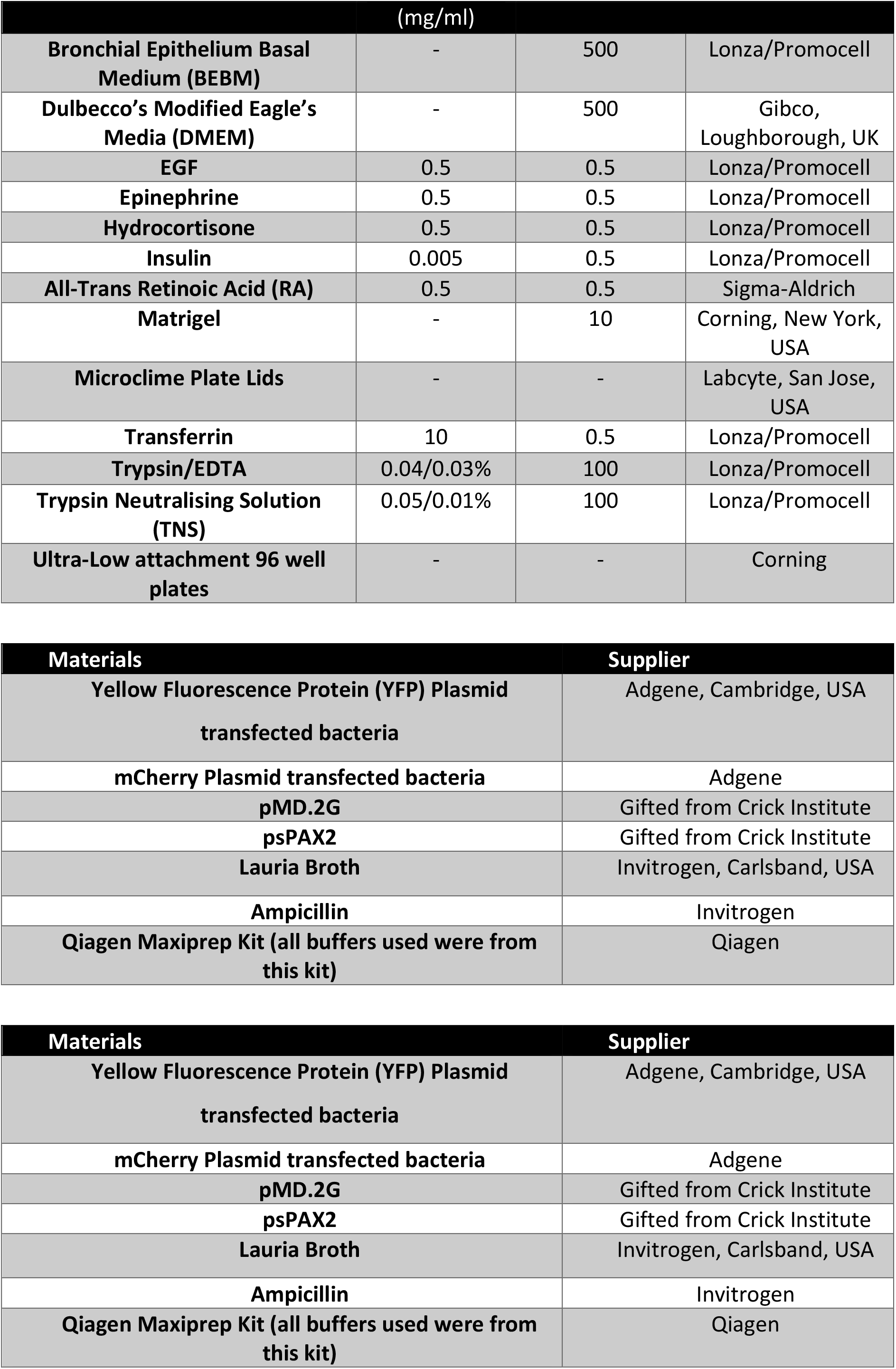
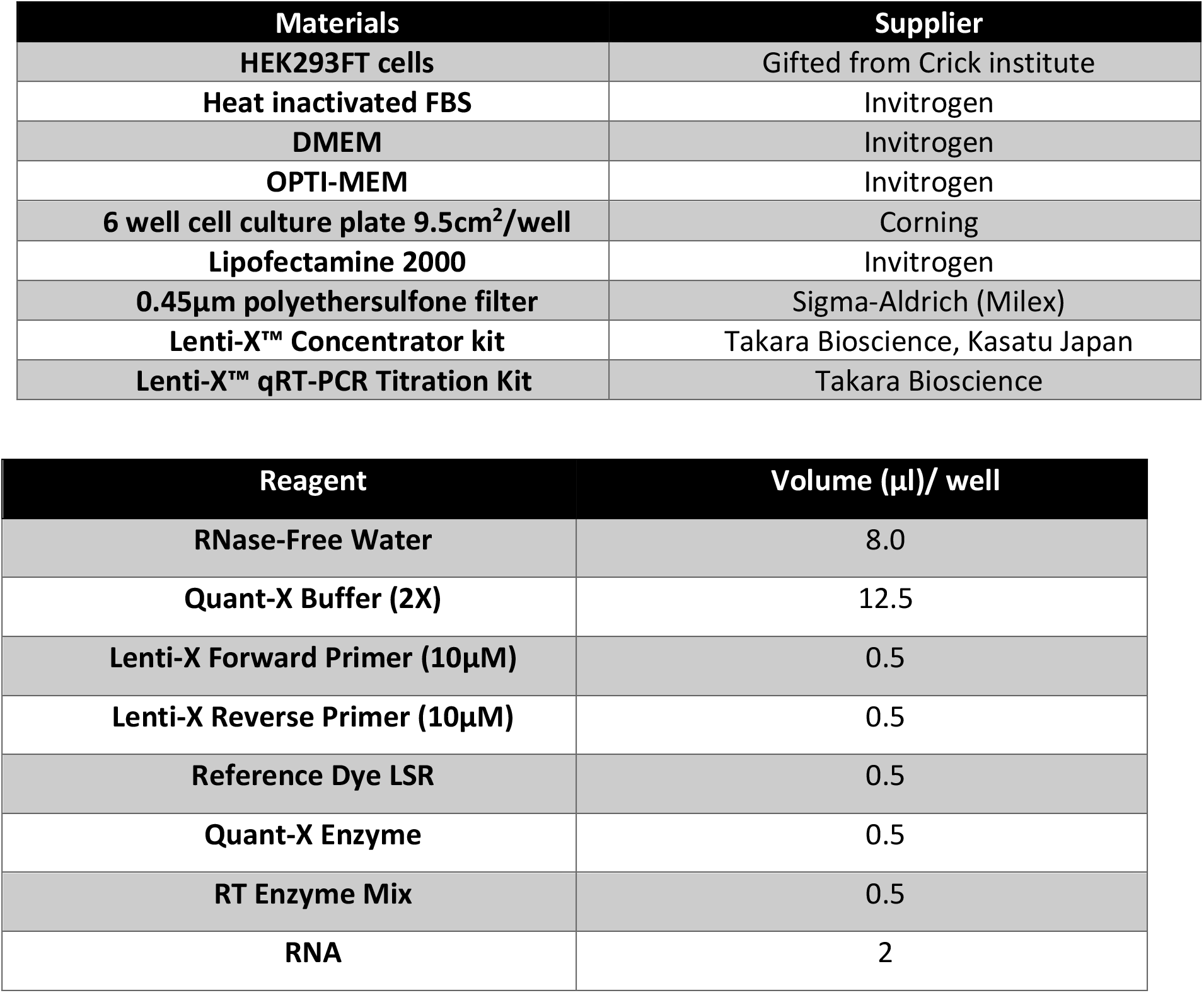

